# “*Haplopappus platylepis* (Asteraceae) resin: an adhesive trap for pest control of crawling arthropods, with antimicrobial potential”

**DOI:** 10.1101/328237

**Authors:** Cristian A. Villagra, Verónica Macias-Marabolí, Constanza Schapheer, Jorge Bórquez, Mario J. Simirgiotis, Javier Echeverría, Marcia González-Teuber, Alejandro Urzúa

## Abstract

The use of plant secondary metabolites has been incorporated as key part of integrated pest management and as an alternative to the use of pesticides. This may even be more relevant regarding domiciliary pest insects, capable of vectoring pathogens to humans. In these environments control its more difficult due to its possible effect on non-target organisms and human health. Here we evaluated the use of the resinous exudate of Chile’s endemic bush *Haplopappus platylepis* (Asteraceae) as a sticky trap for crawling pest insects. We used *Blatta orientalis* Linneus (oriental cockroach), a cosmopolitan synanthropic pest, as test organism. We compared effectiveness on cockroach-trapping of *H. platylepis*’ resin versus a commercially available sticky trap, and analyzed these two sticky substances using UHPLC-DAD-MS and GC-MS. We found that *H. platylepis* resin was as effective as the commercial adhesive on trapping *B. orientalis*. Plant resinous exudate was composed by a mixture of flavonoids, labdane diterpenoids and unsatured fatty acids oxylipins, which are known for their antimicrobial and antioxidant properties. In contrast, the commercial sticky trap was rich in 1-bromohexadecane and 2-clorociclohexanol, which have been described as allergens and as potentially toxic to humans. Considering these findings, we suggest the use of the resinous extract of *H. platylepis* as an effective adhesive trapping method against pest cockroaches and possibly other crawling synanthropic arthropods cohabiting with humans. We highlight the importance of novel, non-toxic and eco-friendly products as strategies to be applied in the management of insect pests.

## Introduction

Synthetic insecticides are controversial as they may represent a potential risk for human health and non-target organisms, beside its contribution to air and soil pollution [1–3]. Furthermore, controlling effects on pests can be rapidly ameliorated due to the evolution of resistance on target organisms [4–6]. This is especially concerning in the case of synanthropic arthropods related to vector-borne and zoonotic diseases inhabiting household, food storage facilities and hospitals [7,8]. These pests are hard to control due to their proximity to human-used spaces, restricting even more the use of various chemical control methods [9,10].

This is the case of several crawling pest arthropods including arachnids such as ticks [11,12], and insects belonging to: Hemiptera, like bedbugs [13] and triatomines [14] and Blattodea: such as pest cockroaches [15,16]. Synanthropic cockroaches [17] such as *Periplaneta americana* (Blattidae), *Blattella germanica* (Ectobiidae) and *Blatta orientalis* (Blattidae) have evolved associated to human-modified environments and usually act as vectors of allergens and diverse pathogenous microorganisms responsible for human diseases [18–21]. Thus, these insects represent a serious threat for human health [22].

The use of insecticides for the control of these insects has been extremely difficult, as cockroaches may become resistant to commonly-used chemical compounds [6]. Moreover, many insecticides at sublethal doses, are repellent to cockroaches and they are capable to avoid its contact [23]. In addition, some studies have shown that the use of pesticide against cockroach infestation paradoxically increases the level of the cockroach allergens Bla g 1 and Bla g 2, and possibly other allergens [24,25]. For example, adults of *B. germanica* exposed to sub-lethal doses of the pesticide boric acid increase the production of the major allergen of Bla g 2 [25], which can lead to significant health problems, including asthma, eczemas skin reactions and allergic rhinitis [26]. Furthermore, it has been demonstrated the evolution of antibiotic resistance in pathogenic strains carried by *P. americana* and *B. germanica* collected from domiciliary and intensive care hospital facilities [27–30].

Therefore, in order to avoid the development of resistances either in the animal or their microbial counterparts, control strategies must combine the suppression of both crawling arthropod vectors and its associated pathogens. This approach must also consider current concerns on the safe use of pesticides for controlling difficult insect pests, especially regarding inhabited and food storaging places [31,32]. In this work we studied the chemical composition of the resinous exudate of a Chilean endemic shrub *Haplopappus platylepis* Phil. (Asteraceae), focusing with particular interest on the presence of antimicrobial potential compounds. Coupled with this, we studied if adhesive extracts of this secretion can be used for the control of pest crawling arthropods, testing its adhesive function against the cosmopolitan pest cockroach *Blatta orientalis* Linnaeus, 1758 (Blattodea: Blattidae).

The use of plant-derived substances, capable of repelling and/or killing synanthropic pests, has been shown in several studies as an effective alternative to insecticides [33–35]. Among these, plant resins have demonstrated to be effective not only against several arthropods [36], but also in the combat against pathogenic microorganisms [37,38]. Moreover, the use of sticky traps could represent a more restrictible pesticide format in comparison with air-borne product, where spray drift unwanted consequences on human health have been reported [39].

In addition, adhesive traps can be displayed in refuge areas where airborne products can not easily reach [40], and reduce pest insects mechanically by catching them [41]. Moreover, these collected insects allow pest density monitoring [42]. This latter is a guide during decision-making for the most appropriate control measurement [43]. Considering the above-mentioned information, adhesive plant secretions such as resinous extractions may arise as suitable candidate for safe pest control of house pest and zoonotic vector insect [44].

*Haplopappus platylepis*, also known as “Devil’s Lollipop”, produces an adhesive resinous secretion covering its leaves and forming a natural sticky trap over floral buds [45]. This plant belongs to an asteraceous lineage presenting copious resin production with known antibacterial and antifungal properties, widely distributed in north and central Chile [38,46–48]. Previously, under field conditions, we showed that *H. platylepis*’ sticky exudate was capable of trapping several groups of insects that were fatally adhered during its blooming season [45]. In this study, we evaluated the potential use of *H. platylepis* inflorescence’s sticky exudate as an alternative adhesive trap for pest crawling insects. For these propose we tested it, in laboratory bioassays, on a common global household pest: the oriental cockroach *B. orientalis*. We compared its effectiveness on adhering pest cockroaches in relation to a commercial adhesive trap (Eco-opción^®^). In addition, we analyzed and compared the chemical composition of the sticky exudate of *H. platylepis* and the commercial adhesive trap using UHPLC-DAD-MS (ultra-high-performance liquid chromatography-diode array detector- mass spectrometry) and GC-MS (gas chromatography-mass spectrometry). Finally, we reviewed for bioactivity of compounds detected in both natural and commercial adhesives, in order to assess both their potential toxicity and harmful effects for humans, as well as any additional biological properties, especially focusing against pathogenic microorganisms.

## Materials and methods

### Plant material and trap extractions

Plant specimens of *Haplopappus platylepis* Phil. (Asteraceae) were determined following Klingenberg’s monography for *Haplopappus* genus [49]. Floral buds of devil’s lollypop were collected during March 2016 at Los Molles, Provincia de Petorca, V Region de Valparaíso, Chile (32°14’07.0"S71°31’24"W) and at Punta Hueso, Pichidangui, Provincia de Choapa, IV Region de Coquimbo, Chile (32°10’27"S 71°31’21"W). Samples were preserved until analysis at −10° C. Voucher specimens (SGO 166498) were deposited in the Herbarium of the “Museo Nacional de Historia Natural” (MCCN), Santiago, Chile.

The sticky exudate of *H. platylepis* was obtained by dipping fresh plant material (300 g) in cold CH_2_Cl_2_ (8 L) for 48 h, following Urzúa 2004’s method[50]. The resulting extract was filtered through a cotton layer and concentrated to a sticky residue (36 g, 12%) Commercial adhesive trap used was Eco-Opción^®^ (Anasac Corporation, Santiago, Chile), sticky trap offered for the control of cursorial domiciliary pest such as ants, cockroaches and spiders. Each unit brings four 29.6×23.3 cm cardboard sticky traps with a total adhesive surface of 11×13 cm. The adhesive mixture from the cardboard was removed with a spatula and followed above-mentioned procedure for extraction. Extracts of both natural (*H. platylepis* inflorescence’s resin) and commercial sticky traps were kept under 4°C for further chemical analyses (see below).

### Insects

Oriental cockroaches used in this work were obtained from a population maintained in our laboratory since year 2014. Further specimens used for this study were collected from locations in San Miguel, Santiago, Metropolitan Region, Chile (33°29’54"S70°38’42"W). For taxonomic identification a general key for cosmopolitan and pest cockroaches present in Chile was used [51]. Insects were kept in captivity under laboratory conditions (20°-25°C and 40%-50% humidity) in 120×50×15 cm plastic rearing boxes, fed with dog food (MasterDog Adult ^®^) and water *ad libitum*, at Instituto de Entomología, UMCE. *Blatta orientalis* from both sexes were used for sticky-trapping bioassays (with body lengths among 5 to 25 mm, measured dorsally from head to last abdominal segments).

### Trapping bioassays

Two treatments and one control were defined for the experiment. Treatments corresponded to cardboard surfaces (40×13cm) painted either with *H. platylepis* resinous exudate or with the commercial trap’s adhesive. For control, a cardboard surface (40×13cm) with no adhesive mixture added was used. Each of these options was presented individually in the experimental arena. For this, the cardboard section was placed in the center of the horizontal space inside the arena, fixing its position with double-contact tape (Fig. 1). For each replicate 10 individuals from different sizes (measured as explained above) were placed in the experimental arena habituation area (Fig. 1), a subdivision of the box from where insect were released without contact them directly. For each trial we lifted the opening section of the habituation area and gave light pulses (10s) during three instances of the experiment: 0, 180 and 360s. At each of these pulses cockroaches tended to leave the habituation area and run to the other extreme of the box crossing the cardboard section. Total time of each test was 6min. After this period, for each treatment and control the number of individuals found attached to the cardboard was counted. Trapped insects were ultimately sacrificed by applying cold temperature (−10 °C). For each of these alternatives we repeated this test 10 times. Before using the experimental arena for each trial, this was cleaned with ethanol (95%), distilled water and dried in order to remove any chemical cue. The response variable was the proportion of insects trapped in each trial for each treatment. As data did not meet the criterion of normality distribution (Hammer, 1999), it was analyzed with a non-parametric analysis of variance Kruskal-Wallis followed by *post hoc* Mann Whitney test. In order to determine if *H. platylepis* inflorescence’s resin and the commercial sticky trap are equally efficient trapping cockroaches of different sizes (seven ranges: from 5 to 7; 8 to 10; 11 to 13; 14 to 16; 17 a 19; 20 to 22 and 23 to 25mm), insect proportion per range, captured in both traps, was compared. This was analyzed by using a Chi square test for two proportions [53]. All analyses were done with the PAST Paleontological Statistic, version 3.15.

**Fig. 1.**
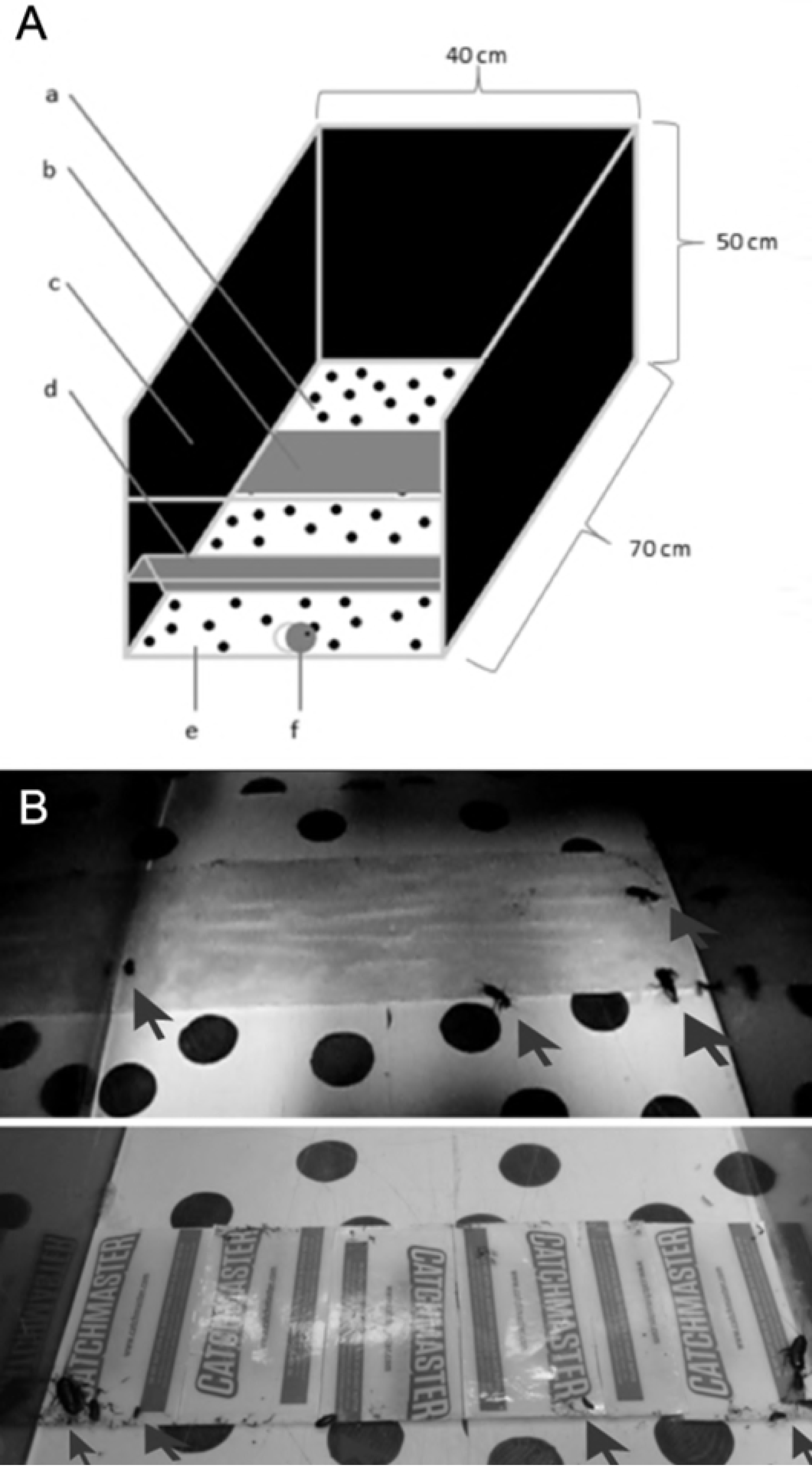
Bioassay setup. **A**. Experimental arena: a. Background pattern. b. Treatment Area, c. Darkened walls, d: Hatch e: Habituation cubicle. f: Arena’s door. **B**. Sticky trap made with *H. platylepis* resin (upper picture) and Eco-opción^®^ adhesive (lower picture). Trapped roaches are highlighted with arrows.

### Chemicals

UHPLC-MS solvents, LC-MS formic acid and reagent grade chloroform were from Merck (Santiago, Chile). Ultrapure water was obtained from a Millipore water purification system (Milli-Q Merck Millipore, Chile). HPLC standards, (kaempferol, quercetin, isorhamnetin, eriodictyol, luteolin, apigenin, naringenin, all standards with purity higher than 95 % by HPLC) were purchased either from Sigma Aldrich (Saint Louis, Mo, USA), ChromaDex (Santa Ana, CA, USA), or Extrasynthèse (Genay, France).

### UHPLC-DAD-MS analyses

Chemical resinous components were analyzed by using ultra-high-performance liquid chromatography-diode array detector-tandem mass spectrometry (UHPLC-DAD-MS). UHPLC-DAD-MS analysis was performed using a Thermo Scientific Dionex Ultimate 3000 UHPLC system hyphenated with a Thermo Q exactive focus machine as it was reported by Simirgiotis et al. (2016). 5 mg of the resinous exudate were dissolved in 2 mL of methanol and filtered with a PTFE filter for a final injection of 10 μL into the instrument. Measurements were done as previously reported by Simirgiotis et al. (2016). The generation of molecular formulas was performed using high resolution accurate mass analysis (HRAM) and matching with the isotopic pattern. Lastly, analyses were confirmed using MS/MS data and comparing the fragments found with the literature.

### LC and MS parameters

Liquid chromatography was performed using an UHPLC C18 column (Acclaim, 150 mm × 4.6 mm ID, 2.5 μm, Thermo Fisher Scientific, Bremen, Germany) operated at 25 °C. The detection wavelengths were 254, 280, 330 and 354 nm, and DAD was recorded from 200 to 800 nm for peak characterization. Mobile phases were 1 % formic aqueous solution (A) and acetonitrile (B). The gradient program time (min, % B) was: (0.00, 5); (5.00, 5); (10.00, 30); (15.00, 30); (20.00, 70); (25.00, 70); (35.00, 5) and 12 minutes for column equilibration before each injection. The flow rate was 1.00 mL min^−1^, and the injection volume was 10 μL. Standards and the resin extract dissolved in methanol were kept at 10°C during storage in the autosampler. The HESI II and Orbitrap spectrometer parameters were optimized as previously reported [54].

### GC-MS analyses

Chemical composition of the commercial adhesive trap was analyzed by gas chromatography-mass spectrometry (GC-MS). GC-MS analysis was performed using a Thermo Scientific Trace GC Ultra linked to an ISQ quadrupole mass spectrometric detector with an integrated data system (Xcalibur 2.0, Thermo Fisher Scientific Inc., Waltham, MA, USA), equipped with a capillary column (Rtx-5 MS, film thickness 0.25 μm, 60 × 0.25 mm, Restek Corporation, Bellefonte, PA, USA) The operating conditions were as follows: on-column injection; injector temperature, 250 °C; detector temperature, 280 °C; carrier gas, He at 1.25 mL/min; oven temperature program: 40 °C increase to 260 °C at 4 °C/min, and then 260 °C for 5 min. The mass spectra were obtained at an ionization voltage of 70 eV. Recording conditions employed a scan time of 1.5 s and a mass range of 40 to 400 amu. The identification of compounds in the chromatographic profiles was achieved by comparison of their mass spectra with a library database (NIST08, NIST, Gaithersburg, MD, USA) and by comparison of their calculated retention indices with those reported in the literature [55] for the same type of column.

## Results

### Trapping bioassays

The proportion of insects found over the cardboards was statistically different among treatments (H (*X*^2^) = 19.43, p < 0.001, Kruskal-Wallis, Fig. 2A). *H. platylepis* inflorescence*’s* sticky exudate and the commercial sticky trap differed with statistical significance from control clean cardboard (in both cases: U Mann-Whitney pairwise, p < 0.001). However, no differences were found in post hoc test for the total number of insects attached on cardboards between the *H. platylepis’* resin and the commercial sticky trap (U Mann-Whitney pairwise, p = 0.691). When the proportion of cockroaches trapped by *H. platylepis’* sticky exudate and by the commercial sticky trap for each size range was compared, no statistical differences were found between natural and commercial sticky traps (*X*^2^ = 1.57, p = 0.211) (Fig. 2B).

**Fig. 2.**
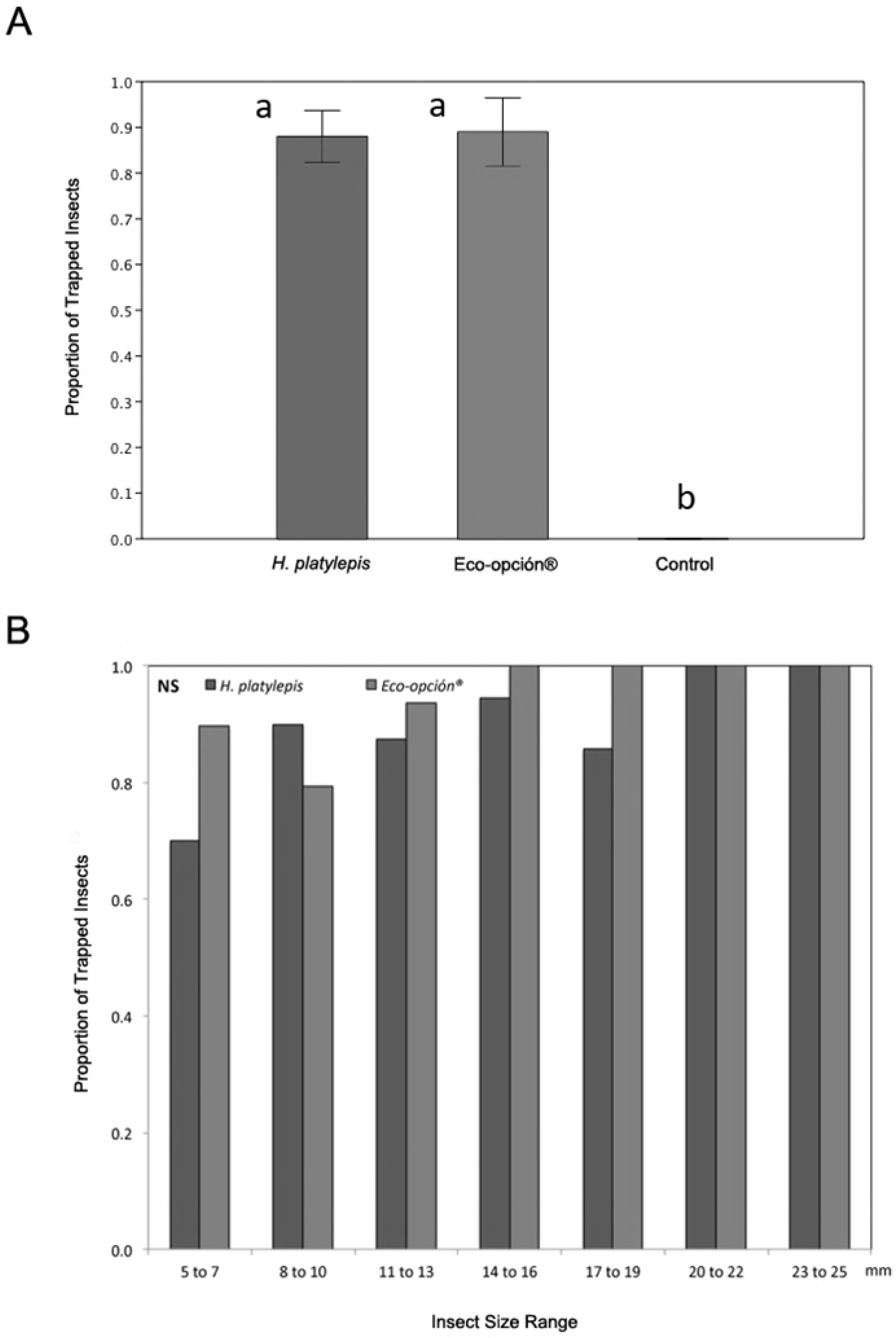
Cockroach adhesion results. A. Mean and 1SE for the proportion of *B. orientalis* found over the cardboard (Y axis) painted with: *H. platylepis* resin (green), Eco-opción^®^ commercial adhesive (red) and control (clean cardboard, black) obtained from 10 replicates each (X axis). Different letters correspond to statistical differences after post hoc test at *p* < 0,05. B. Proportion of cockroaches trapped (Y axis) by either *H. platylepis* resin (light grey) or Eco-opción^®^ commercial adhesive (dark grey) for each insect size range (X axis). No statistical differences were found for each pair compared.

### Chemical analyses

The data-dependent scan experiment was very useful for the identification of unknown compounds since it provides high resolution and accurate mass product ion spectra from precursor ions that are unknown beforehand within a single run. Combining data-dependent scans and MS^n^ experiments, phytochemicals were tentatively identified in *H. platylepis* including simple phenolic acids flavones, flavanones, fatty acids, and labdane diterpenoids. UHPLC Q-orbitrap mass spectrometry analysis of *H. platylepis* sticky exudate showed the presence of twenty seven metabolites in the chromatograms (Fig. 3) including: 7 flavonoids (peaks **5**, **6**, **8-10**, **15** and **16**), 3 phenolic acids (peaks **1-3**), 8 fatty acids (Peaks **4, 7, 13, 14, 18, 21, 22** and **25**), and 9 labdane terpenoids (peaks **11, 12, 17, 19,20, 23, 24, 26,** and **27**). The detailed identification is explained below (Table 1, Figs. 4 and 1S).

**Fig. 3:**
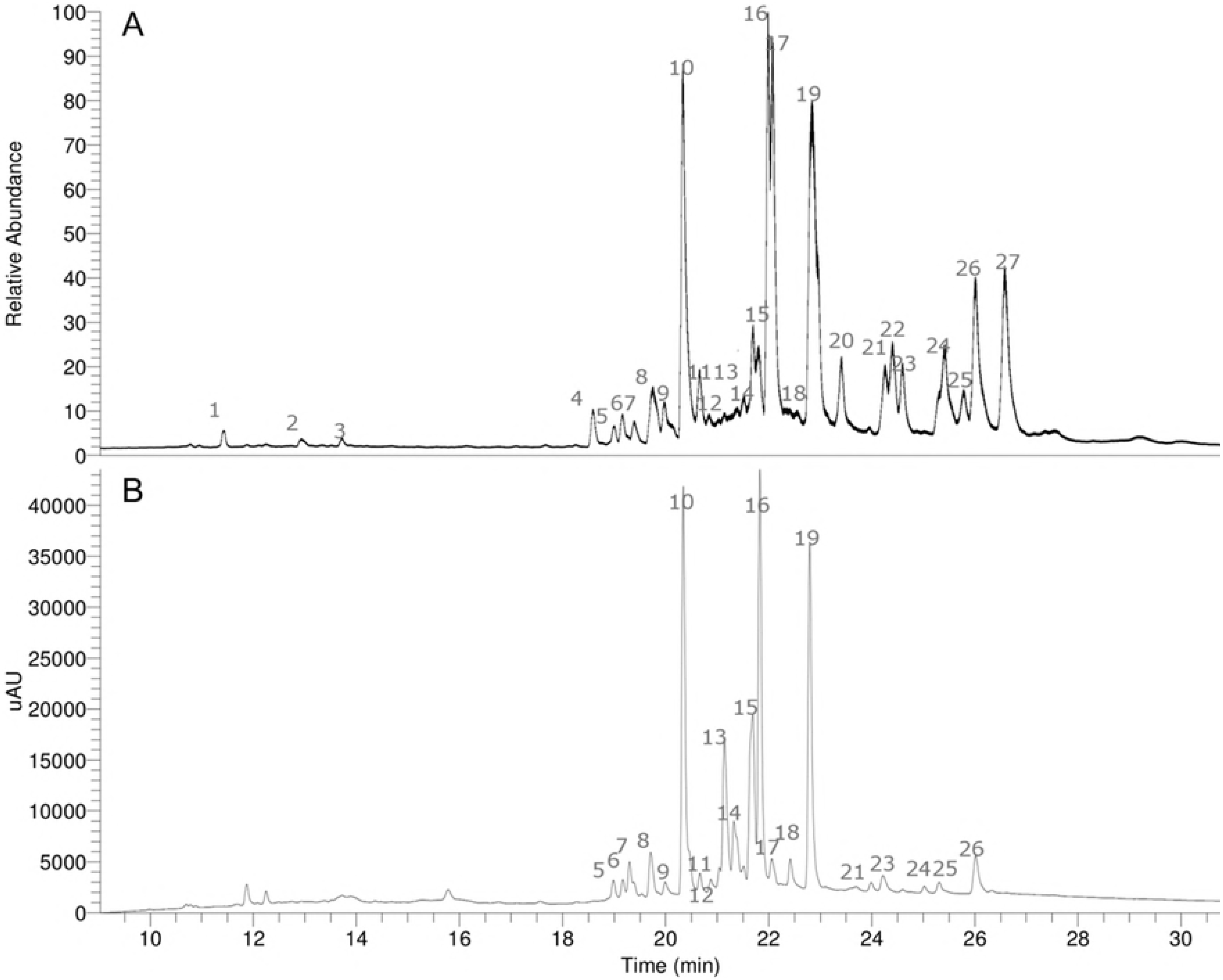
UHPLC chromatograms. A. TIC (total ion current, negative mode) and B. UV at 280 nm, of *H. platylepis* resin.

**Fig. 4:**
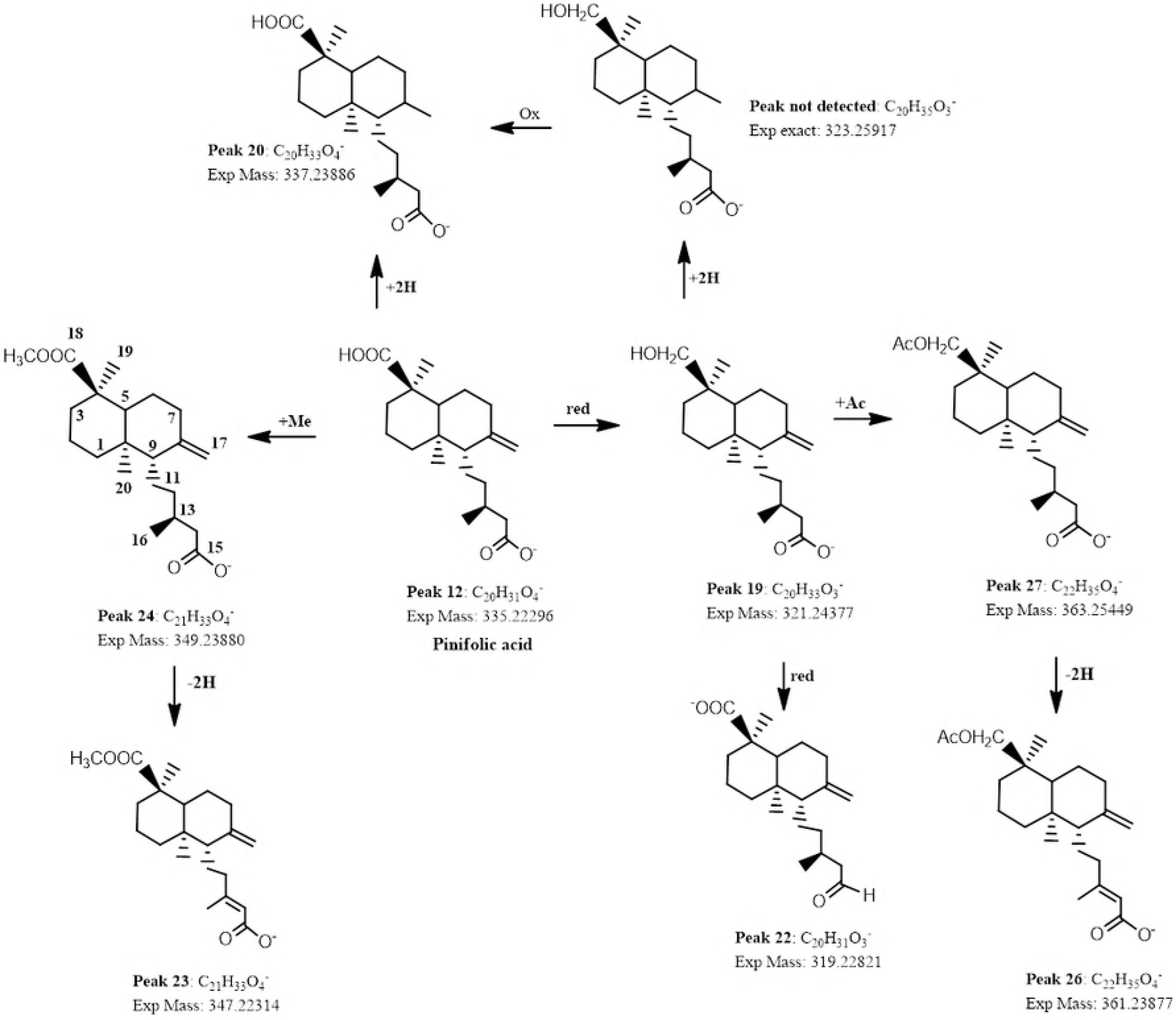
Proposed biogenetic relationships between labdane diterpenoids.

**Table 1.**
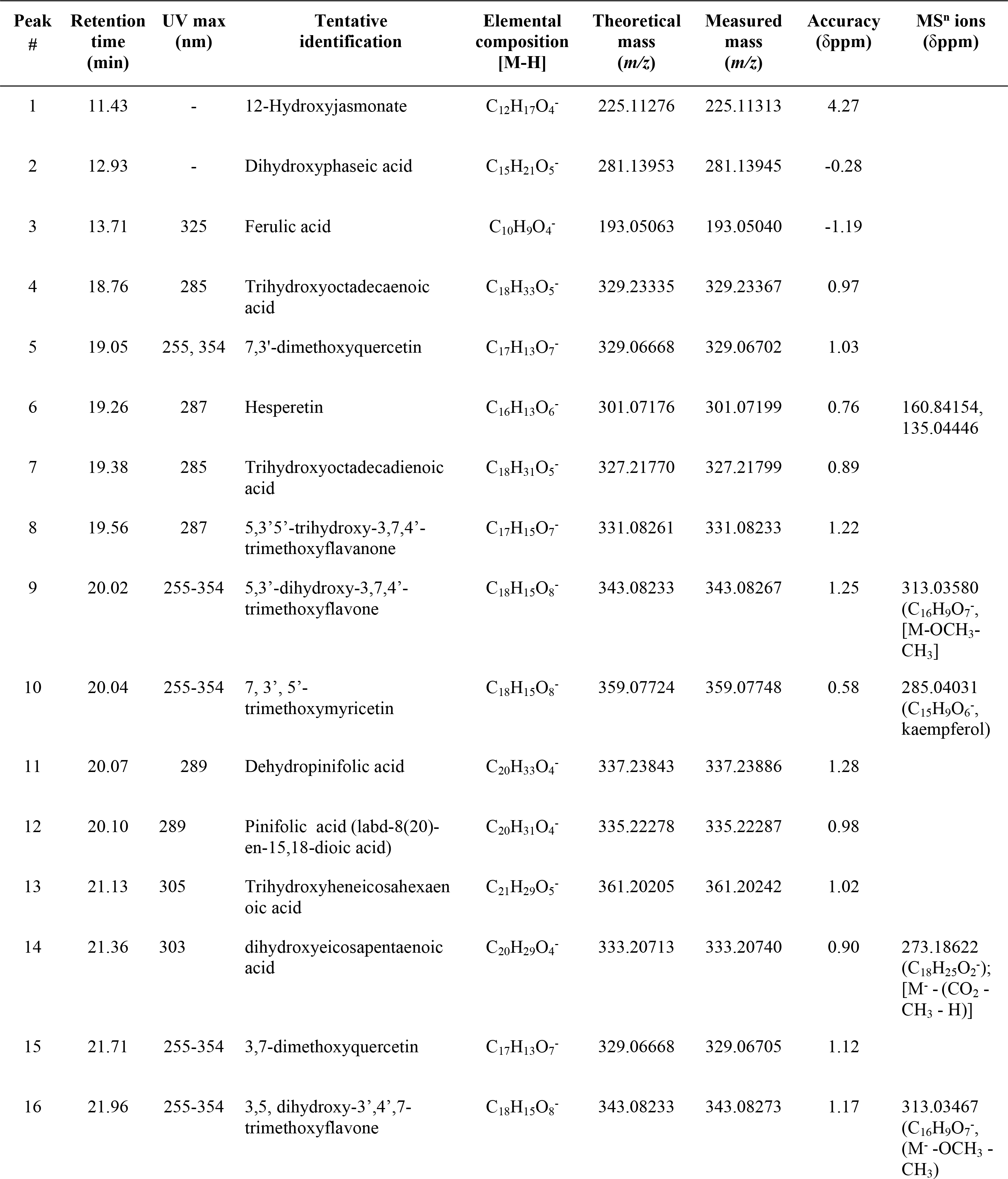
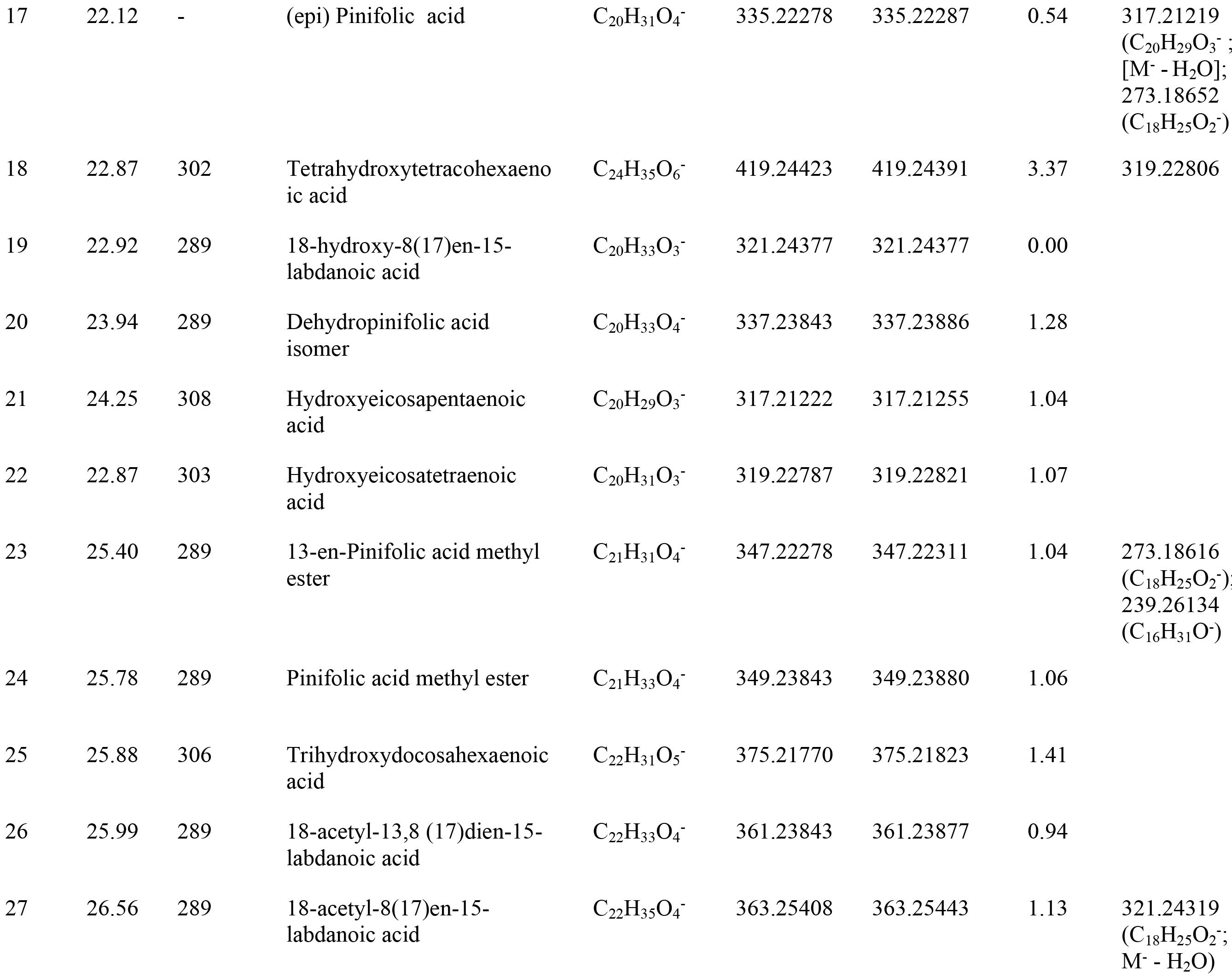
High resolution UHPLC PDA-Q-orbitrap identification of metabolites in *Haplopappus platylepis* resin.

### Flavonoids

Peak **15** with a [M-H]^−^ ion at *m/z* 329.06705 was identified 3,7-dimethoxyquercetin (C_17_H_13_O_7_−) and peak **5** with an ion at *m/z* 329.06702 as its isomer: 7,3’-dimethoxyquercetin (Table 1). Peak **9** with a [M-H]^−^ ion at *m/z* 343.08276 was identified as the trimethoxylated flavonoid 5,3’-dihydroxy-3,7,4’-trimethoxyflavone (C_18_H_15_O_8_^−^), while peak **10** with a [M-H]- ion at m/z 359.07745 as 7,3’,5’-trimethoxymyricetin (C_18_H1_5_O_8_^−^). Peak **16** with a pseudomolecular ion at m/z 343.08273 was identified as 3,5-dihydroxy-3’,4’,7-trimethoxyflavone (C_18_H_15_O_8_^−^). The flavanone hesperetin, peak **6**, have been previously reported as main component in extracts of several *Nolana* species by some of us (Simirgiotis, et al., 2015) and its HR-MS (C_16_H_13_O_6_-) and UV data matched the one obtained in our chromatograms (*m/z*: 301.07176). Another flavanone, peak **8** with a [M-H]^−^ ion at *m/z* 331.08261 was identified as 5,3’,5’-trihydroxy-3,7,4’-trimethoxyflavanone (C_17_H_15_O_7_^−^).

### Phenolic acids

The examination of the chromatograms revealed the presence of 3 phenolic acids: dihydroxyphaseic acid (peak **2**, ion at *m/z* 281.13945, C_15_H_21_O_5_^−^) [56], ferulic acid (peak **1**, *m/z* 193.05040) and 12-hydroxy jasmonate (peak **3**, *m/z* 225.11313) [57].

### Fatty acids

Several peaks were tentatively identified as the dietary antioxidant polyhydroxylated unsaturated fatty acids known as oxylipins [58,59], antioxidant fatty acids. Peak **4** with a [M-H]^−^ ion at *m/z* 329.23367 was identified as trihydroxy-octadecenoic acid (C_18_H_33_O_5_^−^), and peak **7** as its diene derivative (C_18_H_31_O_5_^−^), as previously reported by some of us from Keule fruits [59]. Peak **13** with a pseudomolecular ion at *m/z* 361.20242 was identified as trihydroxyheneicosahexaenoic acid (C_21_H_29_O_5_^−^). Peak **14** with a [M-H]^−^ ion at *m/z* 333.20743 was identified as a dihydroxyeicosapentaenoic acid (C_20_H_29_O_4_^−^) while peak **18** with a [M-H]^−^ ion at *m/z* 419.24391 was identified as dihydroxytetracosatrienoic acid (C_24_H_35_O_6_^−^) [58]. Peak **21** and **22** were identified as hydroxyeicosapentaenoic acid and hydroxyeicosatetraenoic acid (C_20_H_29_O_3_^−^) and (C_20_H_31_O_3_^−^), respectively. Finally, peak **25** with a [M-H]^−^ ion at *m/z* 375.21823 was identified as trihydroxydocosahexaenoic acid (C_22_H_31_O_5_^−^).

### Labdane terpenoids

Labdane terpenoids corresponded to derivatives of pinifolic acid (labd-8(20)-en-15,18-dioic acid, peak **12**, C_20_H_36_O_3_) [60] most of them reported for the first time in this species. Thus, peak **11** with a [M-H]^−^ ion at *m/z* 337.23886 was identified as its hydrogenated derivative of dehydropinifolic acid (C_20_H_33_O_4_^−^) and peak **17** with a [M-H]^−^ ion at *m/z* 335.22296 as an isomer of pinifolic acid (C_20_H_31_O_4_^−^), probably the epimer at C-4 of the latter. Peak **24** was identified as pinifolic acid methyl ester (C_21_H_33_O_4_^−^) and peak **23** as its derivative 13-en-pinifolic acid methyl ester (C_21_H_31_O_4_^−^). Peak **20** with a [M-H]^−^ ion at *m/z* 337,23886 was identified as pinifolic acid derivative (C_20_H_33_O_4_^−^). Three compounds were identified as labdanoic acid derivatives [61]. Thus, peak **19** with a [M-H]^−^ ion at *m/z* 321.24377 was identified as 18-hydroxy-8(17)en-15-labdanoic acid (C_20_H_33_O_3_^−^), Peak **27** with a [M-H]^−^ ion at *m/z* 363.25449 was identified as 18-acetyl-8(17)en-15-labdanoic acid (C_22_H_35_O_4_^−^) and peak **26** as its diene derivative (C_22_H_33_O_4_^−^) (Fig. 4).

### Components identified in the commercial sticky trap

GC-MS identified only two compound in the commercial sticky trap as: 1-bromohexadecane and 2-chlorocyclohexanol.

## Discussion

The aim of this study was to compare the effectiveness of a natural sticky trap against a commercial one in capturing cockroaches by adhesion. In addition, the chemical composition of both traps was analyzed in order to estimate potential harmful effects for humans as well as potential antimicrobial chemical compounds. Our results provide evidence that the natural sticky trap of *H. platylepis* was as effective as the commercial one on trapping pest cockroaches. Considerable differences, however, were found in the chemical composition between the natural and the commercial trap. Whereas the former was rich in plant-derived antimicrobial compounds, the latter was rich in halogenated compounds, whose potential toxic effects for humans have been previously reported.

The *H. platylepis* sticky exudate seems to offer multiple benefits in relation to its use for controlling synanthropic pest crawling insect, such as cockroaches. First, because of its stickiness, it resulted as effective as the commercial trap for capturing cursorial insects, and second, due to its chemical composition rich in antibacterial compounds [62], it shows a further potential for controlling pest arthropod-borne transmitted pathogens. As far as we know, most of the compounds identified for *H. platylepis* resin are reported for the first time in this species. Antibacterial properties of *H. platylepis* sticky exudate can be associated with the phytochemical families detected in the mixture [62]. For instance, flavonoids have shown a wide-sprectrum of inhibitory activity against a variety of human pathogens, including antibiotic-resistant Gram-positive and Gram-negative bacteria, viruses and fungus [62–66]. Labdane diterpenoids are also well known as antimicrobials [67,68]. It has been proved that the presence of a carboxylic acid in the C-15 position, which acted as a hydrogen-bond donor (HBD), is essential for the antibacterial activity of *ent*-labdanes [64]. Furthermore, derivatives of pinifolic acid, which were characterized in the *H. platylepis* sticky exudate, showed this main structural characteristic of labdanes. In addition, pinifolic acid has been previously reported as an effective compound in the treatment of leishmaniasis [69], a global insect-borne disease related to trypanosomes [70]. Long-chain polyunsaturated fatty acids, which were also abundant in *H. platylepis* resin, including oxylipins, have been widely tested for its antimicrobial activity [71–75]. Therefore, further functions of chemical compounds found in *H. platylepis’* resinous exudate expand the potential value of this plant-derived adhesive to act as a control against various vectoring-disease scenarios.

Synanthropic crawling arthropods are usual carriers of several human pathogens [76]. In the case of *B. orientalis*, it has been described to bear several human pathogenic bacteria genera such as *Mycobacteria*, *Klebsiella*, *Staphylococcus*, *Escherichia* and *Enterobacter* [77,78]. Therefore, the occurrence of compounds with anti-microbial functions in the sticky exudate of *H. platylepis* may synergistically contribute as an integrative pest control method, not only directly affecting the insect pests but also its associated pathogenic microorganisms. The commercial sticky trap, in contrast, is poor in its chemical composition and lacks antimicrobial compounds. 1-Bromohexadecane (**1**) and 2-chlorocyclohexanol (**2**) were the only two compounds identified on the commercial trap. Both are known as halogenated compounds. Based on Globally Harmonized System of Classification and Labeling of Chemicals (GHS), both are characterized as irritant for humans, due to the fact that these compounds induce skin corrosion (category 2), respiratory tract irritation (category 3) as well as severe eye irritation (category 2A) (European Chemical Agency-ECHA, 2017). This chemical profile suggests that this commercial trap would not be innocuous for human health; nevertheless, it is commercially offered as an eco-friendly option. Our results highly suggest that *H. platylepis* sticky exudate may be a suitable alternative for controlling synanthropic crawling insects, including cockroaches, at low cost and with additional benefits such as potential antimicrobial properties. These virtues of *H. platylepis* sticky exudate trap fit the current needs and trends in pest control, where several methodologies must be integrated in order to generate novel alternatives in consideration of human and environmental health [79]. Further research is needed in order to test this adhesive resin in other formats for insect trapping as well as to evaluate its effectiveness against other pest insects. For instance, resinous materials have been considered among the updated alternatives for controlling domiciliary termites [44].

## Conclusions

Results here demonstrated that devil’s lollypop resin is a natural source of terpenoids and flavonoids with potential applications as insecticide and antibacterial. Using UHPLC-DAD-MS we have identified 27 secondary metabolites in *H. platylepis’* resin. Most of which, as far as we know, are reported here for the first time. Many of these compounds are flavones, flavanones, phenolic acids, fatty acids, and labdane terpenoids. This chemical knowledge may be helpful for further research on *H. platylepis* and its applications in biomedicine and pest and pathogens control industry. In conclusion, this plant is a rich source of phenolic and clerodane compounds with insecticide and antibacterial activity that may be used as an effective biocontrol agent against zoonotic crawling insects and their associate microorganisms.

## Supporting Information

**Fig. A.1:** Full HR-MS spectra and structures of compounds 3 (a), 9 (b), 10 (c), 12 (d), 14 (e), 22 (f), 23 (g), 26 (h) and 27 (i).

## Acknowledgments

We thank Catherine Cabello and Angel Olguín for help during fieldwork and laboratory work.

## Funding

This research was funded by FONDECYT Iniciación No. 11100109 and CONICYT Inserción No. 79100013 granted to Cristian Villagra, RSG N° 21286-2 to Constanza Schapheer, Proyecto Fortalecimiento USACH USA1799_UA253010, Universidad de Santiago de Chile granted to Alejandro Urzúa, Javier Echeverria and Marcia Gonzalez, and CONICYT PAI/ACADEMIA No. 79160109 to Javier Echeverria.

